# Natural Genetic Variation Shapes Root System Responses to Phytohormones in Arabidopsis

**DOI:** 10.1101/155184

**Authors:** Daniela Ristova, Kristina Metesch, Wolfgang Busch

## Abstract

Plants adjust their architecture by modulating organ growth. This ability is largely dependent on phytohormones. While many genes involved in phytohormone pathways have been identified, it remains unclear to which extent and how these pathways are modulated in non-reference strains and whether this is relevant for local adaptation. Here we assess variation of root traits in response to perturbations of the auxin, cytokinin, and abscisic acid pathways in 192 Arabidopsis accessions. We identify common response patterns, uncover the extent of their modulation by specific genotypes, and find that the Col-0 reference accession is not a good representative of the species in this regard. We conduct GWAS and identify 114 significant associations, most of them relating to ABA treatment. The numerous ABA candidate genes are not enriched for known ABA associated genes indicating that we largely uncovered unknown players. We then study two associated regions in detail and identify the *CRF3* gene as a modulator of multiple hormone pathways. Finally, we show that natural variation in root traits is significantly associated with climate parameters relevant for local adaption in *Arabidopsis thaliana* and that, in particular, ABA regulated lateral root traits are likely to be relevant for adaptation to soil moisture.

**Author Summary:** The root system is a key component for plant survival and productivity. Apart from anchoring the plant, its architecture determines which parts of the soil are foraged for water and nutrients, and it serves as an interface for interaction with microbes and other soil organisms. Plant hormones play crucial roles in the development of root system architecture and its plasticity. However, while there is substantial natural variation of root architectures, it is not clear to which extent genetic variation in hormone related pathways contribute. Using the model species *Arabidopsis thaliana* we quantitatively explore the breadth of natural variation in plant hormone responses to three major plant hormones: auxin, cytokinin, and abscisic acid. The Col-0 reference strain can be quite different from a large proportion of the natural accessions of the species, illustrating a severe caveat of relying on a single reference strain in model species and drawing generalizations from it. Using GWAS, we further identify a large number of loci underlying the variation of responses to plant hormones, in particular to abscisic acid, find links between local adaptation and root responses to hormones, and finally using mutants for GWAS candidate genes, identify novel players involved in regulating hormone responses.

## INTRODUCTION

To grow and survive plants need access to sunlight, nutrients and water resources that are not evenly distributed in the environment. As sessile organisms plants adjust their architecture according to the distribution of resources by modulating organ growth and development. Consequently, plant architecture is of major adaptive relevance (Ackerly et al., 2000). Directed plant growth responses, as well as the resulting plant architecture, are largely dependent on phytohormones, systemic signals that are interpreted in a cellular context (Malamy, 2005). Work related to the phytohormone auxin in the model species *Arabidopsis thaliana* has shown that this pathway is subject to extensive natural variation at the transcriptional level (Delker et al., 2010). However, it is unclear how this relates to plant architecture phenotypes, how the numerous other phytohormone pathways are affected by natural variation, and what the genetic bases for these alterations are. Moreover, it is not known whether the large number of phenotypic and associated molecular responses to phytohormones described in the reference accession Col-0 are representative for *Arabidopsis thaliana* as a species.

The root is an excellent system for studying the dependence of plant architecture on phytohormone pathways. Not only is it technically feasible to perform hormonal perturbations on a large number of roots, but root traits are also of high adaptive relevance as the root system represents the backbone for plant growth and productivity. It anchors the plant to the soil, uptakes water and nutrients, and interacts with soil microorganisms (Den Herder et al., 2010, Lynch, 1995). There are three key developmental processes that determine the most important properties of the spatial distribution of roots, known as root system architecture (RSA): (i) the rate of proliferation and differentiation, which are the two main processes shaping how quickly roots grow; (ii) root growth direction, which determines in which direction the root system is expanding; (iii) the formation of lateral roots, which determine the lateral extensiveness of the root system. All of these processes are under the control of multiple phytohormone pathways (for an overview see (Satbhai et al., 2015)). These hormonal pathways play a key role in adjusting RSA in response to environmental conditions, including availability of the three main macronutrients: nitrogen (Gifford et al., 2013, Krouk et al., 2010, Rosas et al., 2013, Ruffel et al., 2011, Vidal et al., 2010), phosphate (Lopez-Bucio et al., 2005, Nacry et al., 2005, Perez-Torres et al., 2008, Singh et al., 2014), and potassium (Vicente-Agullo et al., 2004) as well as stresses such as salt (Ding et al., 2015, Kumar & Verslues, 2015, Liu et al., 2015, Zhao et al., 2011), drought (Du et al., 2012, Hong et al., 2013, Kang et al., 2002), and exposure to excess metals (Hu et al., 2013, Sun et al., 2010, Yuan et al., 2013).

Here we assess natural variation of RSA traits in response to perturbations of the auxin (IAA), cytokinin (CK), and abscisic acid (ABA) phytohormone pathways by generating and analyzing a comprehensive atlas of these responses in a large set of accessions covering most of the genetic diversity in *Arabidopsis thaliana*. We identify common response patterns and linked RSA traits that are dominated by specific phytohormone pathways, and uncover the extent of their modulation by specific genotypes. Using expression analysis we show that the expression of key genes for multiple hormone pathways are frequently altered in accessions with contrasting RSA responses to hormones. We further study two regions of the genome that are significantly associated with responses in hormone treatments and identify the *CRF3* gene as a being a modulator of multiple hormone pathways. Finally, we show a potential relevance of natural variation of root traits for local adaptation, as we identify significant overlaps between associations for the root traits in this study and environmental variables at the sites of origin of the accessions

## RESULTS

### The Response to Perturbation of Phytohormone Pathways is Subject to Extensive Natural Variation

In order to study and catalogue natural variation of phenotypic responses to phytohormones, we grew a diverse panel of 192 Arabidopsis accessions covering most of the Arabidopsis genetic diversity (Supplemental Figure 1, Supplemental Table 1) on vertical plates for 7 days on 1/5 strength MS medium. We then carefully transferred individual seedlings to different plates supplemented with auxin (IAA), cytokinin (CK), abscisic acid (ABA), or no hormone (C) (Figure 1, Supplemental Figure 2A). Plates were scanned directly after the transfer (day 7) and then again 3 days later (day 10). These images were used to quantify ten root traits (Supplemental Figure 2B and 2C). Overall, we quantified 5 seedlings of 192 accessions in 4 different conditions (3 hormone treatments plus control), totaling 3840 seedlings and 38400 root measurements. There was a very broad spectrum of responses of root growth and development upon perturbation of the endogenous hormone pathways (Figure 1, Supplemental Figure 3). Importantly, these responses were clearly dependent on the genotype, as the broad-sense heritability of the traits was high with an average of 57% for all traits and conditions, ranging from 22% for lateral root density in the branching zone under control conditions to 77% for the primary root length on cytokinin treatment (Supplemental Table 2). In our treatment conditions, we observed strong effects of IAA and ABA on most of the traits, while the impact of CK was rather subtle (Supplemental Figure 3). IAA had strong negative effects on root growth rate (P2) and strong positive effects on lateral root related traits (R, LR.No, LRD_P, TLRL and LRR). ABA had strong negative effects on lateral root related traits (R, LRL, LR.No, LRD_P, LRD_R, TLRL, and LRR), the opposite effect to IAA (Figure 2B). The strongest effects of CK were a negative effect on root growth (P2 and LRL); although we didn’t explicitly quantify it, we also observed a highly increased stimulation of root hair growth upon CK treatment in most of the accessions, especially on the elongated primary root (Figure 2A). Despite being quite genetically diverse, most accessions followed these general response patterns with only some accessions showing strong deviations with regard to specific hormonal perturbations or traits. Surprisingly, Col-0, the reference accession on which most of the previous studies of hormonal effects had been performed, is among the accessions whose root growth responses are frequently quite different from the bulk of Arabidopsis accessions. On average it is in the upper quartile of the accession distribution and belonged to the 1% most extreme phenotypes for some traits in some conditions (Supplemental Figure 3, Supplemental Table 3). Col-0 is therefore not a good representative of the diverse panel of accessions that we have investigated.

**Figure 1.**
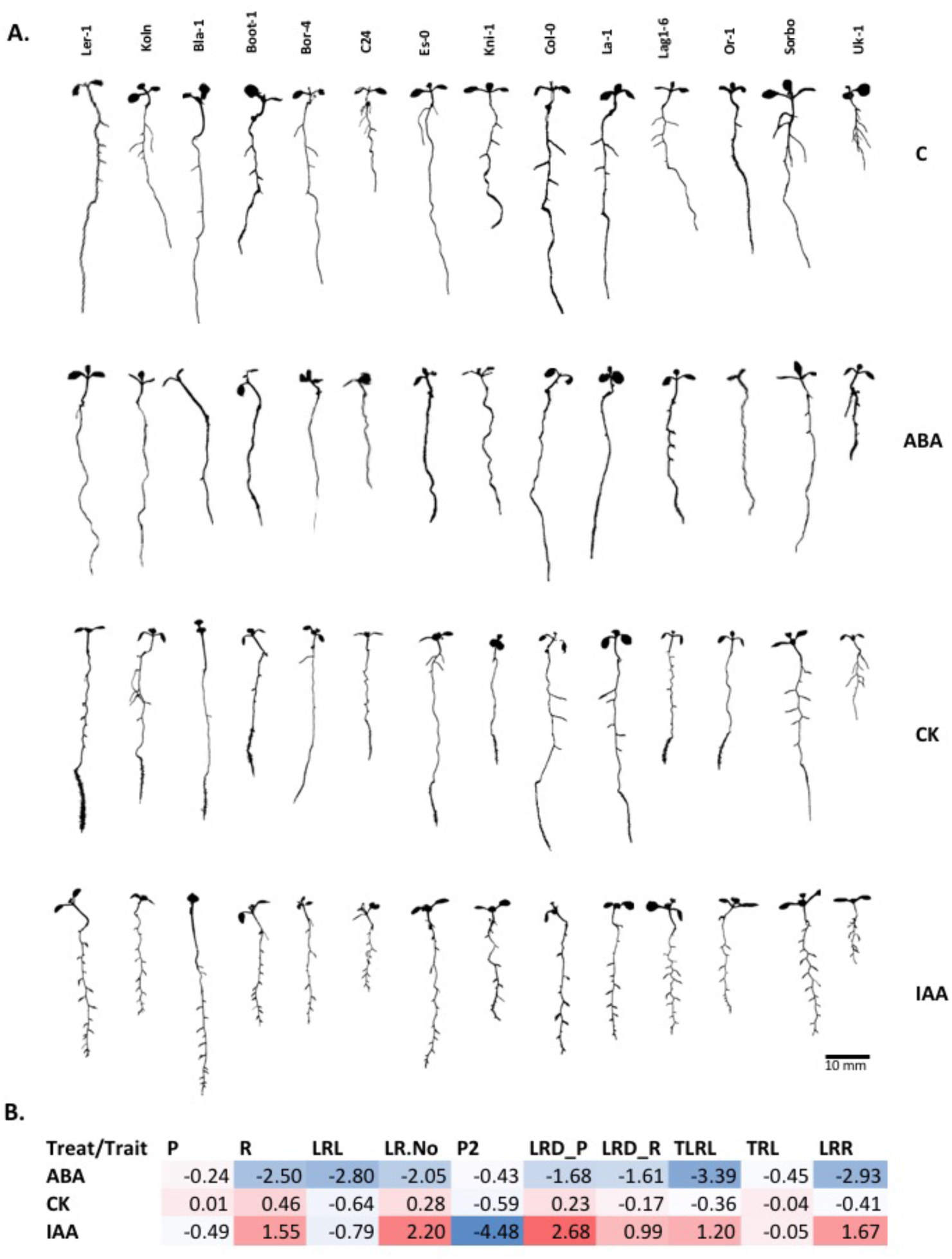
Representative root phenotypes of 14 Arabidopsis accessions on 3 hormone treatments and control. **A)** 192 Arabidopsis accessions, were grown on control conditions for 7 days followed by transfer to media supplemented with IAA, CK, and ABA, or no hormone (C, control). Plates were scanned on day 10, and root traits were quantified and segmented using FIJI. Here we show segmented seedlings of 14 accessions. **B)** Heatmap of (log2) fold-change for each trait and hormone treatment compared to control treatment across all 192 accessions and 4 conditions.

**Figure 2.**
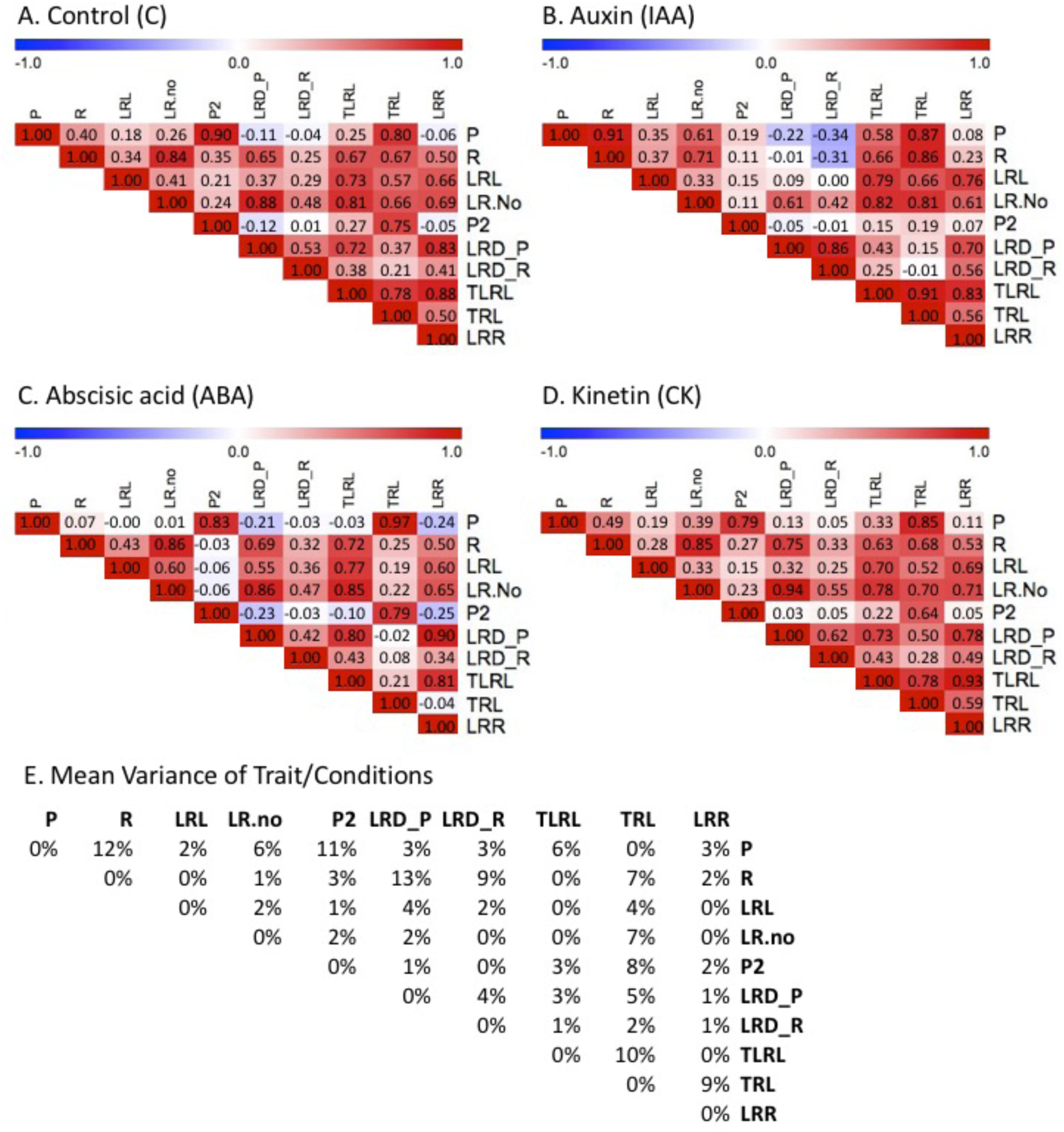
Patterns of Root Trait Correlations. (A-D) Heatmaps of pairwise correlations (Pearson product-moment correlation coefficient) of 10 root traits in 192 accessions upon transfer on control (A), auxin,(B), abscisic acid (C), and cytokinin (D) medium. Colors gradient indicates negative (blue) or positive (red) correlation coefficients. (E) Mean variance of all traits in percent. Traits: primary root length (P), growth of P between day 7 and 10 (P2), branching zone of P (R), average lateral root length (LRL), and lateral root numbers (LR.no). Other traits are calculated by formulas: total LRL, TLRL (LR.L*LR.No), total root length, TRL (TLRL+P), density of P, LRD_P (LR.No/P), density of R, LRD_R (LR.No/R), and length ratio, LRR (TLRL/P).

**Table 1.**
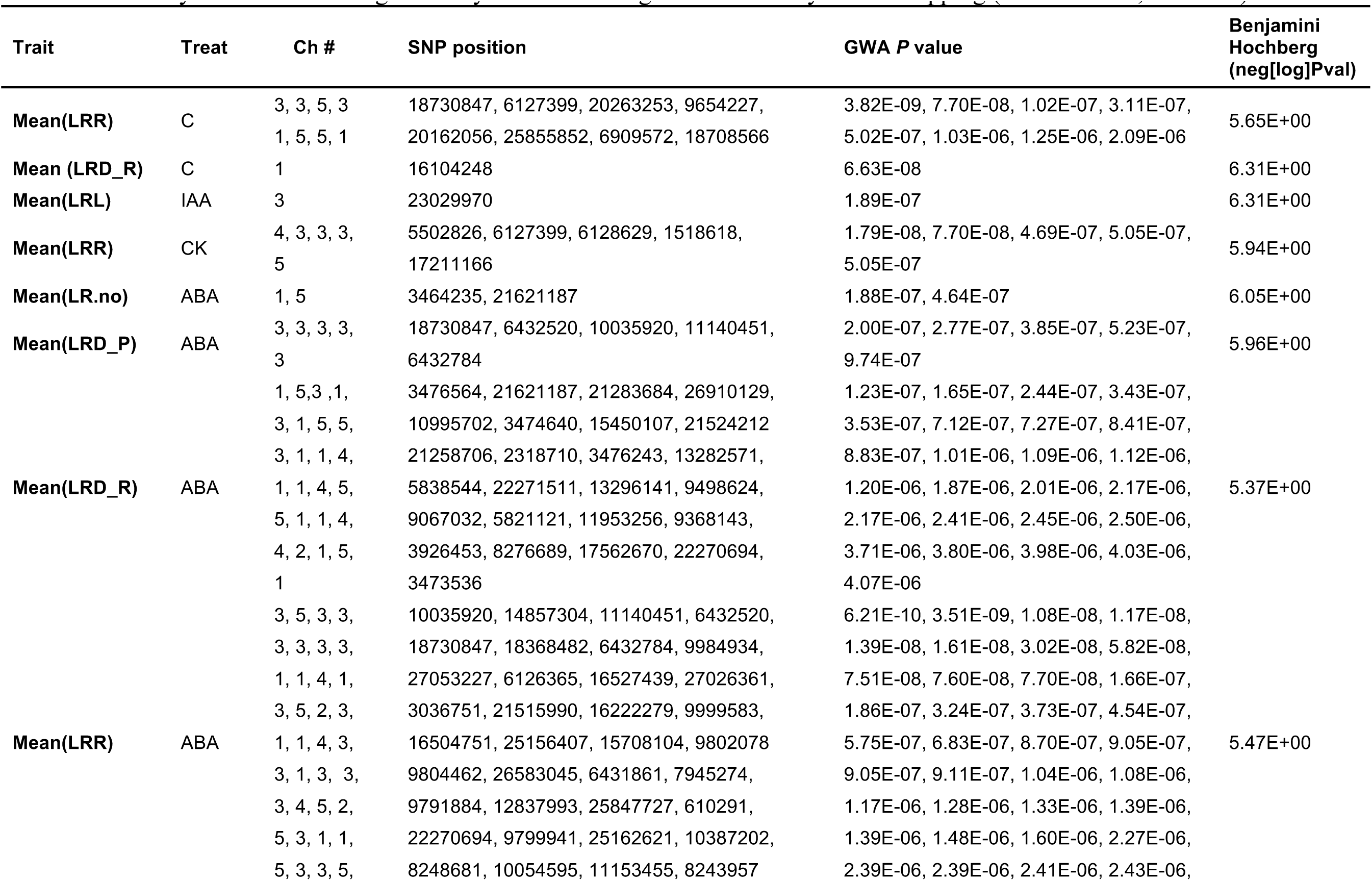
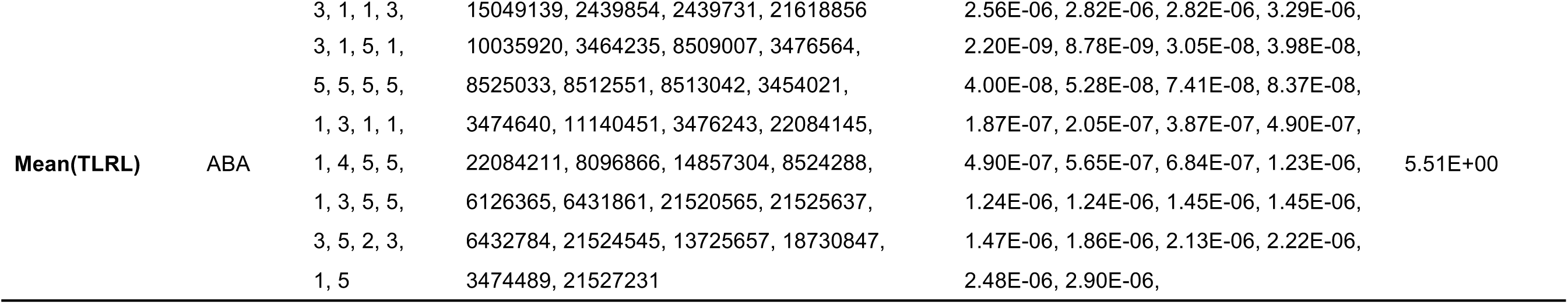
Summary Information of Significantly Associated Regions obtained by GWA Mapping (BH threshold, 5% FDR).

In conclusion, we have generated the first comprehensive atlas of root responses to perturbation of hormonal pathways. Our data show that while there are clear trends of phenotypic responses to perturbation of specific hormonal pathways, there is significant natural variation in these responses. Overall, this suggests that hormonal signaling pathways within the *Arabidopsis thaliana* species are largely acting on the same traits, but that the extent of the hormonal control of specific traits is genotype dependent. We therefore conclude that hormonal pathways are subject to natural genetic variation. Importantly, the effects of hormones on root growth in the Col-0 reference accession are not representative of the major proportion of Arabidopsis accessions.

### Distinct Hormonal Pathways Dominate Distinct Traits

The dependence or independence of traits in diverse genotypes is highly relevant for inferring the genetic architecture of a trait; a strong dependence suggests a common genetic, and possibly molecular, basis underlying the regulation of the respective traits. Our comprehensive atlas of RSA traits of a large-number of Arabidopsis genotypes upon perturbation of phytohormone pathways allowed us to ask which of the hormonally regulated RSA traits were linked and to which extent. We used our trait data from the 192 accessions and calculated pairwise Pearson’s correlation coefficients of all traits for each condition (Figure 2A-D). To determine whether the extent of trait correlation was similar in all conditions, or whether perturbations of phytohormone pathways affected a particular trait correlation, we also calculated the variance of each trait correlation between all conditions. We reasoned that if this variance was high, the observed trait correlation would be strongly dependent on a subset of phytohormone pathways (Figure 2E). The highest variation of correlations between independent traits was observed for the length of branching zone and the lateral root density of the primary root (σ=0.13). While these traits are highly correlated in control (0.67), CK (0.63) and ABA (0.67) conditions, this correlation reverses upon IAA treatment (-0.31). This strong effect can be traced back to the impact of IAA on root growth rate and its impact on the branching zone, which are key for the second and third ranked correlation variances respectively. Under control conditions, there is a very high positive correlation between primary root growth rate before treatment and growth rate after treatment (r=0.9), indicating that root length after 7 days is generally a very good predictor for root length after 10 days. IAA treatment, however, breaks the correlation of these two traits almost completely (r=0.19). ABA and CK treatments diminish this correlation only slightly, suggesting that the observed effect of IAA on root growth rate is highly specific. Overall, this strongly suggests that the regulation of root growth rate is strongly dominated by the auxin pathway, as demonstrated by multiple studies (Evans et al., 1994; Coenen and Lomax, 1997; Tian and Reed, 1999; Friml, 2003; Swarup and Bennett, 2003; Robert and Friml, 2009; De Smet et al., 2012; Pacifici et al., 2015). While primary root length and the length of the branching zone are moderately positively correlated under control conditions (r=0.4), this correlation increases to very high levels under IAA treatment (r=0.91), suggesting that auxin generally promotes lateral root formation or outgrowth. ABA treatment renders these two traits almost completely uncorrelated (r=0.07), suggesting that ABA generally represses the formation or outgrowth of lateral roots, a finding that is consistent with the literature (De Smet et al., 2003).

Overall, these examples not only demonstrate the significance of specific hormone pathways for specific traits, but also support the view that RSA traits such as branching and primary root growth rate can be regulated independently of each other, dependent on the condition, and over a large number of genetic backgrounds, a phenomenon previously shown for the impact of different nitrogen environments on root traits (Gifford et al., 2013).

### Genotypes can Alter Phenotypic Responses to Hormones

While our correlation analysis revealed common trends over a large number of genotypes, our atlas of root responses to perturbation of hormonal pathways also allowed us to study which genotypes modulate the responses to perturbations of hormone pathways. To do so, we performed a hierarchical-clustering of our ten traits separated by condition. Different clusters contain groups of accessions with genotypes that generate a similar RSA under the respective conditions (Figure 3, and Supplemental Table 4). We observed that perturbation of specific hormone pathways shift RSA to a distinct state (for instance, IAA causes roots to become shorter and more branched) but that the genotype determines the extent to which this state is reached. For instance, IAA treatment cluster 5 is shifted to the IAA RSA state to a lesser degree than IAA cluster 6. Overall, our analysis partitions the phenotypic space that we explored using the systematic perturbation of hormone pathways and illustrates the tremendous impact of the genotype in determining how root development responds to hormones. However, while this analysis is highly useful for identifying classes of accessions and visually illustrates the effects of hormones and how these effects differ, it was based on capturing RSA using individual traits, some of which are dependent on each other and some of them not. Moreover, hierarchical clustering gives insight into classes, rather than continuous similarities. To gain a more integrative insight into the action of hormones, we conducted a principal component analysis (PCA) on all our traits for all conditions (10 root traits across 192 accessions and four treatments). 99% of RSA variation (given by our 10 traits) could be captured by five PCs, while 9 PCs are needed to explain the complete variation (Supplemental Table 5). The first PC captures 62% of the total variation. Several traits contribute to it, including root number (LR.No), lateral root density (LRD_P and LRD_R), total lateral root length (TLRL), and length ratio (LRR). The second PC accounts for 26% of the variation and represents mainly primary root length (P) and total root length (TRL) (Supplemental table 5). Comparable contributions of the same root traits to each PC (PC loadings), as well as PC variability were quantified for the same hormonal treatments of the reference accession Col-0 (Ristova et al., 2013), demonstrating a highly reproducible pattern of the RSA traits and their relations to hormonal perturbations. Overall, IAA treatment, and to a lesser extent ABA treatment, results in a clear separation from the other three conditions when plotting the most informative PCs, PC1 and PC2, while CK and C are largely overlapping (Figure 4A). To identify accessions with genotypes that alter from the norm of how RSA responds to hormone pathway perturbation, we calculated the Euclidian distance to the average PC values in the RSA space defined by PC1 and PC2 for each accession and each condition. This distance indicates the deviation of a particular genotype to the average RSA response of all accessions (Supplemental Table 6). For each accession, we then calculated the average Euclidian distance of all conditions (Supplemental Table 6). This average represents a measure of how profoundly a genotype differs from the norm in altering its RSA in response to all hormonal perturbations that we had performed. Eight accessions (Sh-0, Lu-1, Col-0, Sorbo, UKSE06-373, Kz-1, Or-0, Uk-1) were more than 2 standard deviations from the mean of all accessions in this regard (Figure 4A, Supplemental Table 6), indicating that they show significant alterations of their hormonal response from the average *Arabidopsis thaliana* accession. Again, the Col-0 reference accessions was among these accessions, again highlighting the conclusion that much of what has been learned in studies on Col-0 may not represent the species as a whole. Taken together, these data demonstrate that hormone perturbations generally lead to specific patterns of developmental responses that shift RSA from one configuration to another (i.e. IAA shifts RSA to shorter and more branched configurations). However, these response patterns can be significantly altered by genotypic variation. This genotypic variation must therefore be in the pathways that perceive or respond to phytohormone cues. Overall, this demonstrates that natural genetic variation in phytohormone pathways can be a major contributor in shaping the Arabidopsis root system.

**Figure 3.**
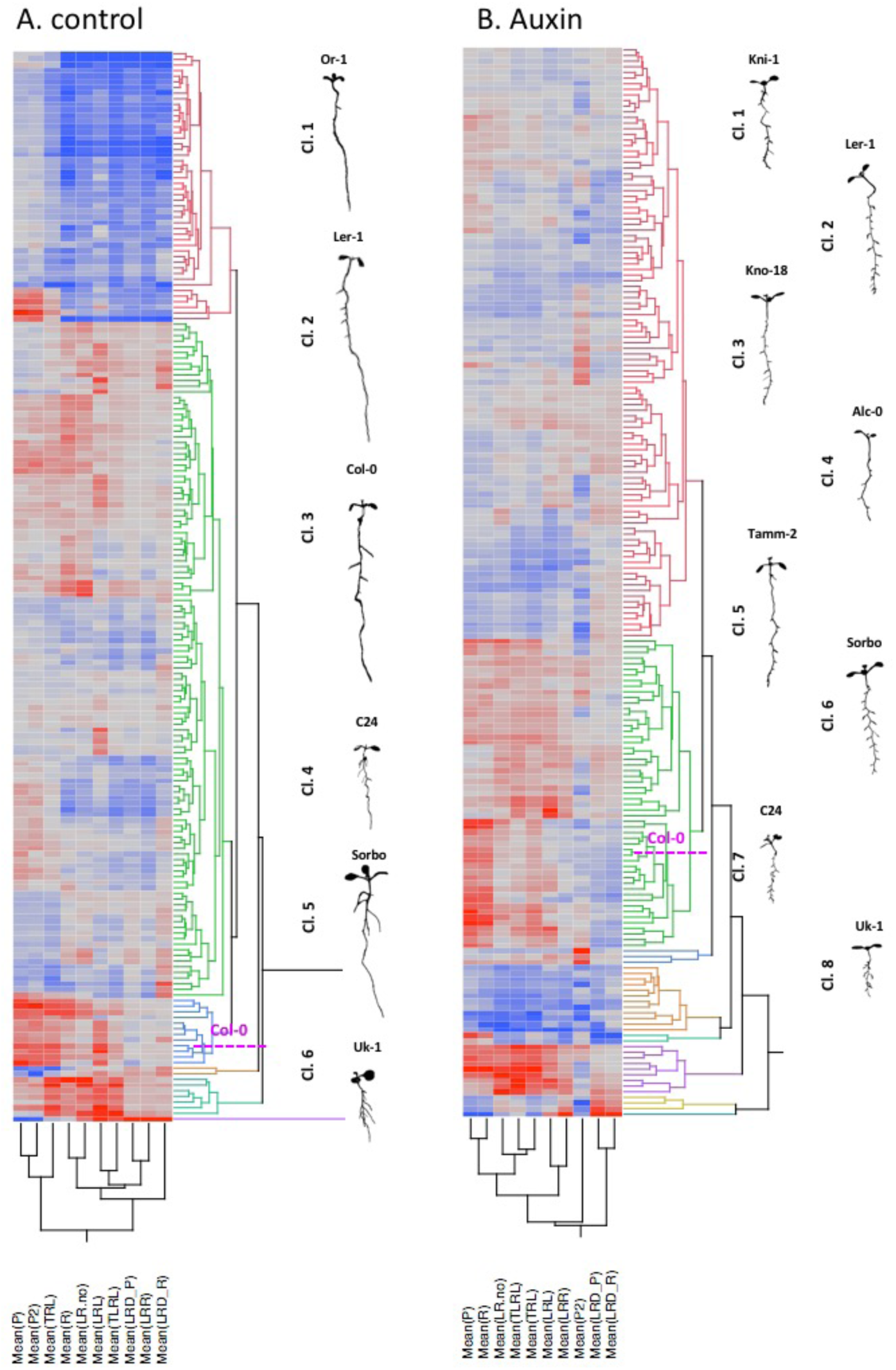
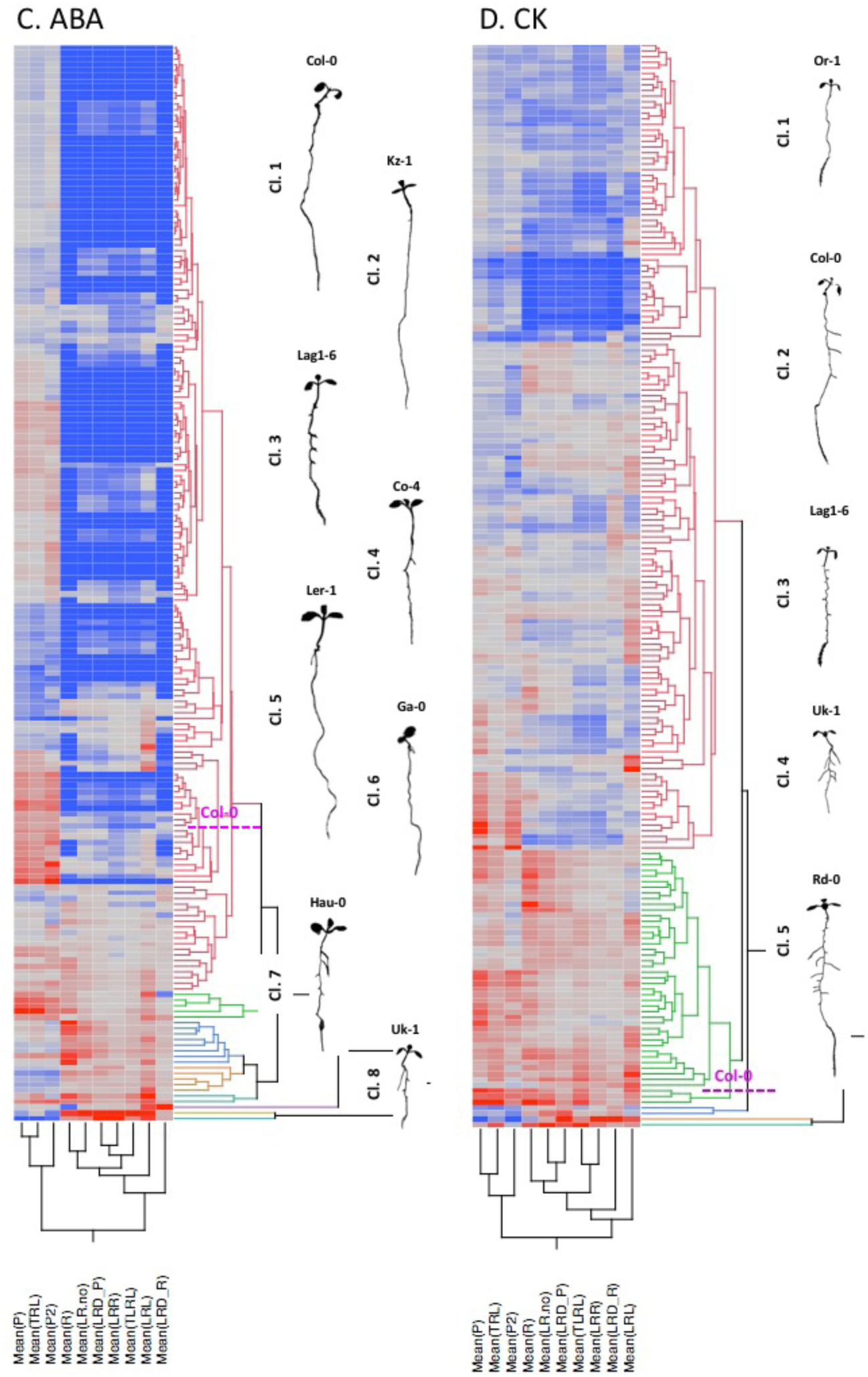
Hierarchical Clustering (HC) of Arabidopsis accessions by Condition. HC was performed on mean root traits of 192 accessions, in each of the three hormones (ABA, CK, and IAA). On the right side of each dendrogram we show one representative accession from each cluster. **A)** HC of 192 accessions on C (control, no hormone added). **B)** HC of 192 accessions on IAA. **C)** HC of 190 accessions on ABA. D). HC of 192 accessions on cytokinin (CK). Cluster numbers and additional info in Supplemental Table 4.

**Figure 4.**
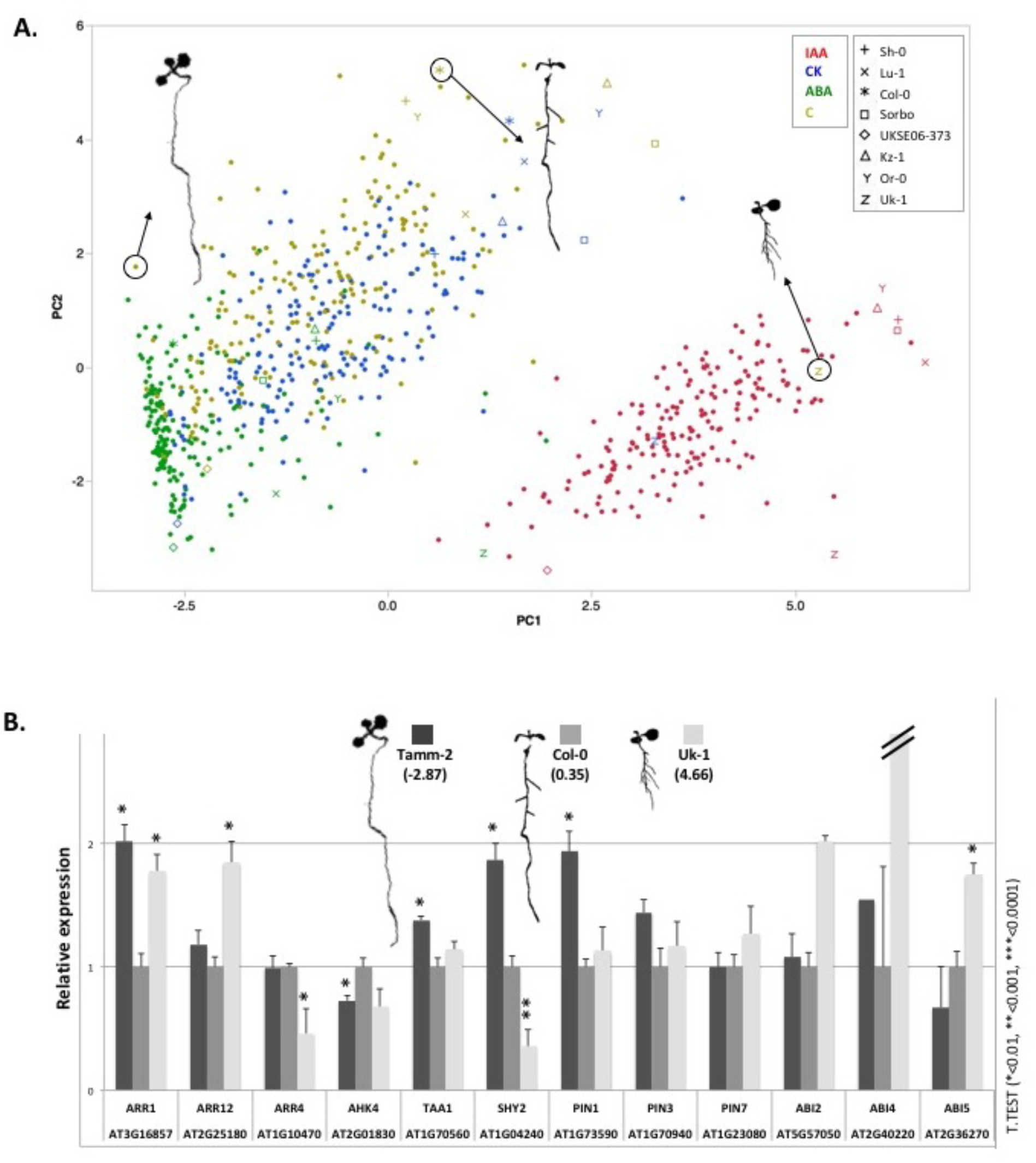
Multivariate Analysis of 192 *Arabidopsis* accessions on 4 conditions and gene expression analysis from extreme accessions. Mean values for 10 root traits across 192 Arabidopsis accessions and 4 conditions (IAA, CK, and ABA, and no hormone C) were used to perform a Principal Component Analysis (PCA). **A)** Biplots of the ten quantified root traits and 192 accessions under 4 conditions are shown for the first 2 principal components (PCs). Each accession in each condition is represented by a dot, and each condition has a different color. Each extreme accession is displayed by a specific symbol that is colored according to the treatment condition. **B)** Relative expression of 12 genes involved in IAA, CK, and ABA hormonal signaling pathways in extreme accessions quantified by PC1: accession Tamm-2 (negative extreme in PC1), accession Uk-1 (positive extreme in PC1), and the reference accession Col-0. Segmented seedling phenotypes of the 3 accessions are shown at the top, including the PC1 value, and the position of each accession in control conditions is circled in A). A Student’s T-test was performed to determine the significance of gene expression changes in the extreme accessions compared to the reference accession. Mean with s.e.m. is shown. Data from 3-4 biological and 2 technical replicates. *: *P* < 0.01

### Genetic Variation leads to Changes in the Molecular Regulation of Hormone Pathways

To test whether the phenotypic differences we observed are reflected at the molecular level, we measured the expression of genes related to hormone signaling in accessions that represented different PC1 values (Uk-1: high PC1; Tamm-2: low PC1; Col-0: intermediate PC1). We included 8 genes of the *SHY2* network which are involved in auxin, cytokinin and abscisic acid signaling, and 4 additional genes involved in auxin biosynthesis (*TAA1*) (Stepanova et al., 2008), abscisic acid signaling (*ABA2*) (Leung et al., 1997), and cytokinin signaling (*ARR4, AHK4*) (Salome et al., 2006, Yamada et al., 2001). We found that 8 of 12 genes were expressed at different levels in these accessions, several of them in a pattern similar to the PC values (Figure 4B). Thus, the steady state expression level of important components of hormone pathways can be vastly different in different accessions. To corroborate this finding, we also made use of a public dataset of root transcriptome data of 7 accessions (Delker et al., 2010). Under control conditions, there was also significant variation in expression of these key genes, providing further evidence the expression levels of genes in hormonal pathways are highly dependent on the genotype (Supplemental Figure 4). Overall, this demonstrates that there can be a wide variety of transcriptional states in hormonal pathways relevant to RSA in different genotypes.

### Genome-wide Association (GWA) Mapping on RSA under Hormone Treatments

Our analysis of root responses to hormone pathway perturbation clearly showed that these responses are under genetic control. To identify genetic variants that were associated with these different responses, we performed genome wide association (GWA) mapping on all traits in all conditions (10 traits * 4 conditions) using a mixed model algorithm (Kang et al., 2008, Yu et al., 2006), that was previously shown to correct for population structure confounding (Seren et al., 2012), and using SNP data from the 250K SNP chip (Atwell et al., 2010, Brachi et al., 2010, Horton et al., 2012). We used a Benjamini Hochberg threshold (5% FDR) to select significant associations. In total we identified 114 significant associations, corresponding to 441 genes within a 20 kb window of the associated SNPs (Table 1, Supplemental table 7). We found associations for each treatment, with ABA perturbation yielding the most at 98; CK had 5 significant associations and IAA only one. GWAS for the control condition without hormone treatment added 9 associations (Table 1, Supplemental table 7). We observed only 3 cases where GWAS identified the same SNP associated with different treatment conditions and the same trait (Supplemental Figure 5).

Despite the large number of significant associations for the ABA pathway, the genes close to these associations didn’t contain obvious core ABA signaling related genes. To test whether this absence of *bona fide* ABA candidate genes was statistically significant, we identified all genes for which a role in the ABA pathway had been assigned based on reported experimental evidence or mutant phenotypes (Supplemental table 8). There was only one gene that overlapped between GWAS-based candidate gene list and the annotation-based ABA *bona fide* candidate gene list. We then tested whether this overlap is expected by chance, we found that our observed overlap was at the lower limit (towards our GWAS list being depleted of *bona fide* candidate genes) of what was to be expected by chance (p<0.16; Supplemental Figure 6). This clearly showed that *bona fide* candidate genes are not overrepresented in our list of GWAS candidates and indicates that, for complex traits such as root responses to hormonal treatment, genes targeted by natural variation are frequently not the genes found by traditional experimental methods (such as forward genetic mutant screens).

To identify high confidence candidate genes whose variants underlie the associated traits, we focused on two cases. The first case involved the trait LRR (ratio of total lateral root length and primary root length) where a significant association on chromosome 3 was found under both control (C) and cytokinin (CK) conditions (Supplemental Figure 7). Approximately 7KB from the most significantly associated SNP (Chromosome 3; 6127399) is a gene with a known role in the hormonal regulation of development, *JASMONATE-INSENSITIVE 3* (*JAI3*, AT3G17860) (Chini et al., 2007). While the top SNP itself was located in the gene body of AT3G17910 (Figure 5B), the entire region surrounding this SNP contained many single-nucleotide polymorphisms in a pattern consistent with the phenotypes of contrasting accessions (Supplemental Figure 7A). When analyzing linkage disequilibrium (LD) in this region, we found that SNPs in only two genes (AT3G17890 and AT3G17900) were in LD (R^2^>0.4) with the SNP that we had identified through GWAS. We tested whether the expression of any gene in this region was correlated with the LRR trait. Only the three genes in LD showed expression levels correlated with the LRR phenotype. Of these three genes, AT3G17910 showed the highest correlation (Figure 5C, Supplemental Figure 7B). *JAI3*, the most likely candidate gene based on its known role, did not show any correlation. To test whether one or more of these genes are involved in regulating lateral root related traits, we obtained available T-DNA insertion mutant lines for AT3G17890 (SALK_050958), AT3G17910 (SALK_045920) and *JAI3* (*jai3-1)* (Chini et al., 2007). No insertion line in the genic region of AT3G17900 was available. The expression levels of the respective genes were significantly reduced in the SALK lines that we obtained (Supplemental Figure 7C). We examined the phenotypes of these three mutants under control and cytokinin treatment conditions. While the *jai3* line showed an overall reduced root growth in both conditions, the response to CK was altered for multiple traits in T-DNA mutants of the other two genes (Figure 5D and 5E). In particular, alterations were observed in both mutants for the branching zone (R) and lateral root number (LR.No) traits, as well as for primary root length (P) and total root length (TRL) in one mutant (SALK_045920). While compared to Col-0, the SALK_045920 (AT3G17910) line showed a decreased branching zone under control and an increase of it upon CK treatment, the second line SALK_050958 (AT3G17890) showed a slight decreased response upon CK treatment (Figure 5D). Moreover, expression levels of several genes in hormonal pathways were significantly altered in the SALK_045920 (AT3G17910) line and SALK_050958 (AT3G17890) (Supplemental figure 7D), with a much more pronounced perturbation of gene expression in the SALK_045920 line. Overall, this strongly suggests that the yet uncharacterized genes AT3G17890 and AT3G17910 are involved in regulating aspects of CK hormonal signaling that are relevant for shaping the same root traits in a complex manner. We note that we can’t evaluate the role of the third gene, AT3G17900, which is also a potential candidate based on the LD pattern and the expression level correlation.

In contrast to the first case, a well-known candidate gene was identified for another trait. Here, a significant SNP (chromosome 5; position 21621187) for lateral root number (LR.No) and root density (LRD_R) (Figure 6A) under ABA treatment was located in the promoter region of *CYTOKININ RESPONSE FACTOR3* (*CRF3*, AT5G53290) (Figure 6A, Supplemental figure 8A). *CRF3* has been shown to be a key link between cytokinin signaling and auxin transport in root development, acting downstream of cytokinin perception and transcriptionally regulating auxin transport (PINs) (Simaskova et al., 2015). As this region was devoid of other genes and a SNP in the coding region of *CRF3* was in LD (R^2^>0.4), this was the only and best candidate gene. To test whether we would obtain phenotypes that consistent with the GWAS phenotypes in our conditions, we obtained a loss of function T-DNA line (SAIL_240_H09) for *CRF3* that has been described previously (Simaskova et al., 2015). We found that this line had an overall reduced root growth, affecting both root length and lateral root number, when grown under control and abscisic acid (ABA) conditions (Figure 6C and 6D). While the most pronounced LR effect upon ABA in the wild type is a strong reduction of the lateral root length (LRL and TLRL), these traits were not affected by ABA in the *crf3* mutant line, leading to a highly significant interaction of the *crf3* mutation and ABA treatment (Figure 6C and 6D). Moreover, expression levels of genes related to auxin, cytokinin and ABA hormonal pathways were significantly altered in the *crf3* line (Supplemental figure 8B). Taken together, this shows that *CRF3* not only links auxin and cytokinin responses, but also impinges on ABA dependent root growth regulation.

**Figure 5.**
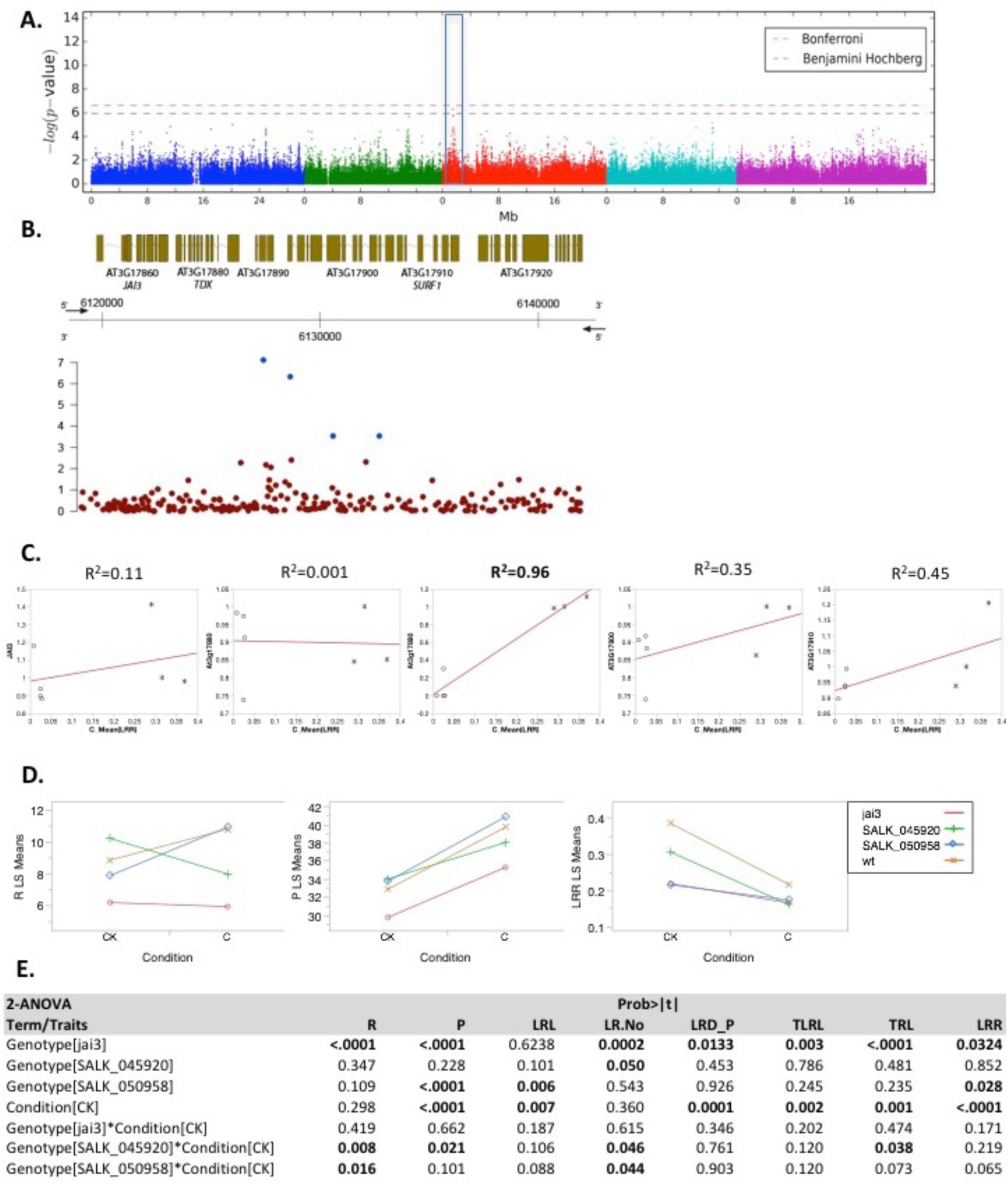
Genome-wide association and mutant analysis. A) Manhattan plot of length ratio, LRR (total length of lateral roots/total primary root length) under CK treatment (a significant association is present for the same trait under control condition). B) The genomic region surrounding the significant GWA peak. Bottom, −log10 *P* values of association of the SNPs. SNPs highlighted in blue are in in LD (R^2^ > 0.4) with top SNP. Top, gene models in genomic regions. The *x* axis represents the position on chromosome 3. C) Regression analysis of the associated trait LRR (mean) and the relative expression of the 5 genes in the associated region in several extreme accessions. D) Reaction norm plots of root branching zone (R, in mm), primary root length (P, in mm), and length ratio (LRR) for available T-DNA lines for genes in the associated region E) 2-way-ANOVA summary for all root traits quantified under control (C) and cytokinin (CK) treatment (P-value=0.05). Mutant lines: *jai3-1* (At3g17860), SALK_050958 (At3g17890), SALK_045920(At3g17910); (N>30).

**Figure 6.**
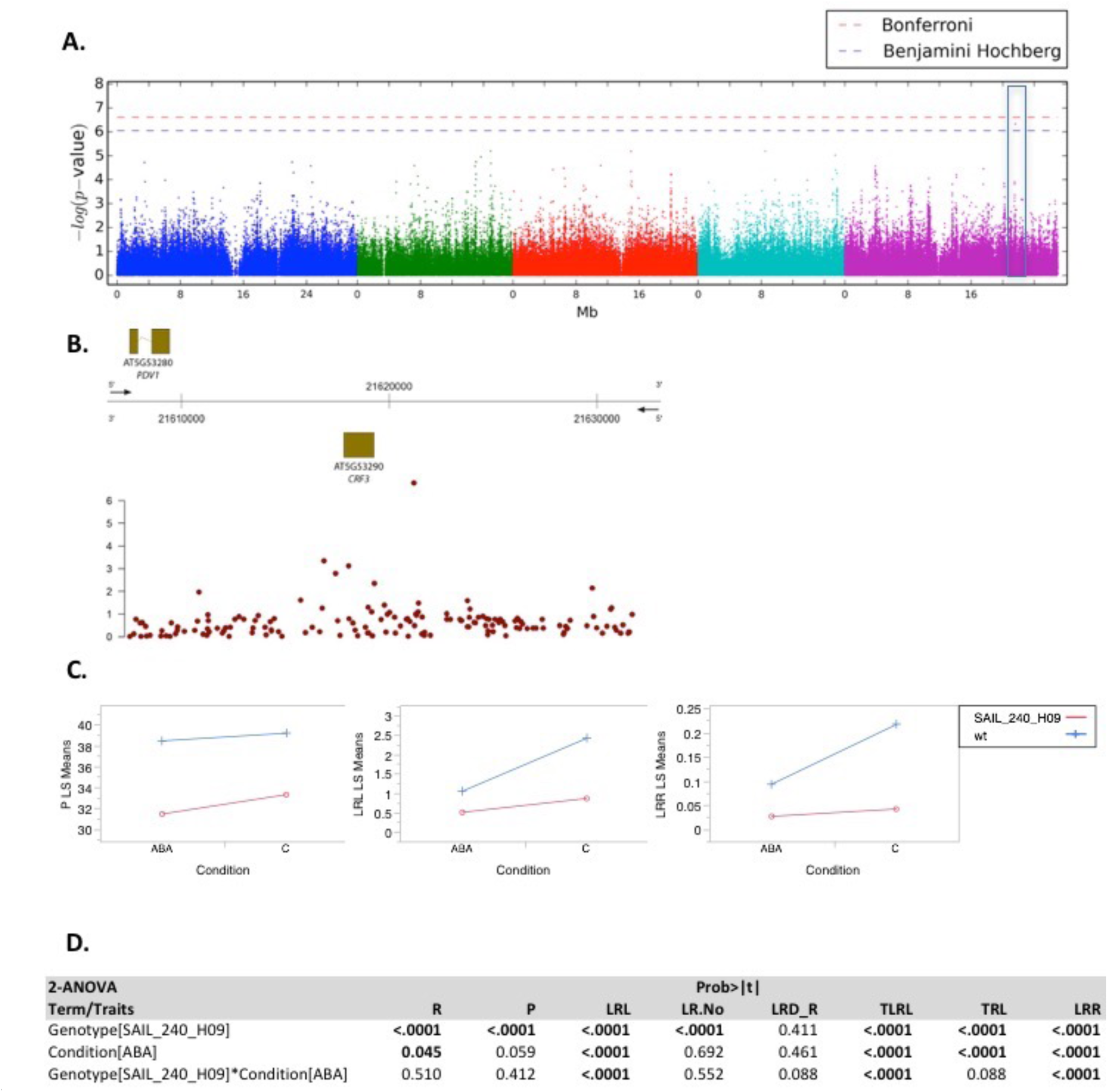
Genome-wide association and mutant analysis of CRF3. A) Manhattan plot of the significant genome-wide association for trait root density (LRD_R) (ch. 5: 21621187). B) The genomic region surrounding the significant GWA peak. Bottom, −log10 *P* values of association of the SNPs. Top, gene models in genomic regions. The *x* axis represents the position on chromosome 5. C) Reaction norm plots of primary root length (P, in mm), lateral root length (LRL, in mm), and length ratio (LRR) for the T-DNA line for CRF3 (SAIL_240_H09), the only gene in the associated region (20 kb window). D) 2-way-ANOVA summary for all root traits quantified under control (C) and abscisic acid (ABA) treatment (N>30).

### Natural Variation in Hormone Signaling Pathways and Local Adaptation

As natural variation in pathways that affect root growth responses to phytohormones has a tremendous impact on root system architecture related traits, it was tempting to speculate that the variation we observed was relevant for local adaptation. Much of the local adaptation in Arabidopsis is driven by climate related parameters (Fournier-Level et al., 2011). We therefore first determined whether there is an overlap between the genetic variants associated with the root traits that we quantified in our study and the genetic variants associated with climate parameters at the site of origin of the accessions. For this, we treated the climate parameters of every accession’s site of origin as a trait, and conducted the same population structure corrected GWAS procedure that we had for our trait data on 35 climate variables capturing biologically relevant climate aspects (Bioclim variables) (Hutchinson et al., 2009, Kriticos et al., 2012) for all accessions of the RegMap panel (Horton et al., 2012). We then determined the overlap top SNPs (0.5 top percentile) between each climate parameter GWAS and each GWAS from our RSA-hormone study and computed the probability of this overlap occurring by chance. After correction for multiple testing, out of the 1400 computed overlaps of phytohormone induced RSA responses associated top SNPs and climate variable associated top SNPs, 15 overlaps showed a significant (FDR < 5%) overlap (Supplemental Table 9). We then tested whether the root traits and the climate traits were significantly correlated when accounting for population structure and correcting for multiple testing. We found that out of these 15 trait combinations that had a significant overlap of associated SNPs, 9 displayed a significant population structure corrected trait correlation (FDR < 5%) and another 2 were marginally significant (FDR < 10%) (Supplemental Table 9). Overall this suggests that the root traits that we measured in our study are involved in local adaptation.

The most significant overlap between SNPs was found between ABA induced lateral root density and “Mean moisture index of coldest quarter” (FDR < 0.2%), consistent with the well-known role of ABA in response to water related responses. The trait correlation of this most overlapping combination was negative (r= -0.189610096), suggesting that less lateral roots are produced in an ABA dependent manner in accessions that are derived from areas in which the soil contains high moisture in the coldest quarter of the year. As this trait correlation is marginally significant when correcting for population structure (FDR < 7.5%), this suggests that variation in genes that determine the impact of ABA on lateral root development is adaptive in environments with contrasting soil moistures. Interestingly, none of the genes in proximity to the 19 overlapping SNPs is a *bona fide* ABA gene (Supplemental Table 10).

While the most significant overlap at the SNP level clearly suggested an important role of ABA for local adaption for soil moisture, auxin and cytokinin treatments were also among the list of root traits that were significantly overlapping with environmental parameters at the level of SNP overlap and at least marginally significant correlated to the same climate trait (FDR < 8%).

However, 7 out of the 11 combinations displaying significant overlaps at the level of SNPs and traits did not associate with hormonal treatment, but with RSA traits under non-treated conditions. Finally, only one of these overlapping traits is related to primary root growth. However, as only 2 out of 10 traits that we had scored related specifically to the primary root, this is still in the realm of expectation. Overall, these data clearly show that natural variation in root traits is significantly associated with parameters that are highly relevant for local adaption in *Arabidopsis thaliana* and that, in particular, ABA regulated lateral root traits seem to be relevant for adaptation to soil moisture.

## DISCUSSION

### Natural Variation of Hormone Signaling Pathways and Root Architecture

In this study, we report the first comprehensive atlas of RSA responses to hormonal perturbations in a large number of *Arabidopsis thaliana* natural accessions. Our results show general patterns of phenotypic responses to the perturbation of specific hormonal pathways, as well as significant natural variation in these responses across a set of accessions that captures a large fraction of the genetic variation in Arabidopsis (Supplemental figure 1). We show that specific hormonal pathways dominate a distinct subset of traits, as perturbations of specific pathways can break correlations that exist between traits under control conditions (Figure 2E). In our experimental conditions, auxin perturbation caused the most dominant effect on RSA by strongly repressing root growth rate and stimulating lateral root growth. ABA treatment reduced primary root length, as well as lateral root numbers (Figure 1B), and diminishing the correlation between these two traits, thereby demonstrating its negative role in the emergence of lateral roots. In contrast to the replication of these well described effects of auxin and abscisic acid (De Smet et al., 2003, Evans et al., 1994), our cytokinin treatment conditions did not lead to the previously observed strong inhibitory effects on primary and lateral root growth (Laplaze et al., 2007, Li et al., 2006). We did however observe the stimulatory effect of CKs on root hair growth which had been described before (Werner et al., 2001). A possible reason for the observed cytokinin response in our experimental conditions might be the specific cytokinin chosen and the concentration of nutrients in the medium, as we used kinetin and one-fifth-strength MS medium.

Regardless of the specifics of our treatments, our atlas of root growth responses revealed the extent of variation in the responses to perturbation of hormone pathways. Our hierarchical clustering results identified groups of accessions sharing similar or diverse responses to a particular hormone perturbation (Figure 3). Importantly, these clusters can be very useful for choosing parents for quantitative trait loci mapping (QTL) via linkage mapping approaches using recombinant inbred lines (RILs). While current mapping populations are based on only a small subset of accessions (Alonso-Blanco & Koornneef, 2000, Koornneef et al., 2004), they have already been used to identify *BREVIS RADIX (BRX)*, a gene that is responsible for the hormone related short-root phenontype of the UK-1 accession (Mouchel et al., 2004, Mouchel et al., 2006). Our results provide an opportunity for choosing novel, previously unexplored accessions with diverse root responses to hormones as parents for creating new populations to identify novel alleles underlying hormone pathway dependent root developmental processes. Principal component analysis (PCA) showed that hormones can shift RSA through the phenotypic space (Figure 4A). In our conditions, auxin treatment led to the most pronounced phenotypic effect on RSA compared to ABA and CK. Of course, we cannot exclude that other concentrations of hormones and their interplay with nutrient concentrations could lead to even more pronounced CK or ABA effects, but our findings corroborate the prominent role of auxin in root development. Moreover, the PCA analysis allowed us to systematically identify outlier accessions that behave clearly differently than the bulk of Arabidopsis accessions.

Importantly, accessions that are different with respect to their RSA response upon hormonal treatments also show deviations in the expression patterns of hormone related genes in control conditions (no hormone treatment). This shows that the varying responses we observed upon hormonal stimuli are associated with different transcriptional states of the hormone signaling networks. This conclusion is consistent with genome-scale observations in 7 accessions before and upon auxin treatment, in which vastly different transcriptome responses were detected (Delker et al., 2010). Importantly, such different transcriptional states that presumably underlie altered responses to hormones can be caused quite indirectly, for instance by alterations of the balance of another hormone pathway. This has been nicely illustrated by studies related to the *BRX* gene that is causal for a large proportion of the root phenotype of the UK-1 accession. There, the lack of auxin response in the *brx* mutant results from a root specific deficiency in brassinosteroid production (Mouchel et al., 2006). Given the strong and widespread intraspecies variation that we have observed here, it is not surprising that distinct differences in the wiring and interaction of plant hormone pathway exist between species (Pacheco-Villalobos et al., 2013)

Taken together our results show that the root growth responses of Arabidopsis accessions upon the perturbation of hormone pathways display distinct patterns that depend on the hormone shifting RSA towards distinct states, and that the extent of this shift depends on genetic factors. This demonstrates that hormonal pathways are subject to natural genetic variation within a species, and it is tempting to speculate that this might contribute to adaptation to diverse soil environments.

### Reference accessions and the species

Much progress in uncovering the genetic and molecular bases of traits has been made in reference strains of model species, including in Arabidopsis, Drosophila, and yeast. Findings in these reference strains have been frequently generalized for their respective species and often even for a whole phylogenetic branch of species. Our data demonstrate that the response of a reference strain can be quite different from a large proportion of the natural populations of the species. Within our diverse panel of 192 Arabidopsis accessions, the reference strain Col-0 belongs to the accessions that are least representative of RSA responses upon perturbation of hormone pathways (Supplemental Table 3, Figure 3 and Figure 4). This is not simply a peculiarity of our investigation; for instance, in a study dissecting 107 diverse phenotypes (Atwell et al., 2010), Col-0 is located in the 1% lower tail of the distribution of all accessions for 14% of the traits (Supplemental Figure 11). While these findings don’t affect the validity of the fundamental mechanisms for hormone responses that have mainly been discovered in Col-0, they suggest that, at a certain level of detail, studies will uncover relations and mechanisms that are only found in specific genotypes within a species. We note that we don’t suggest that Col-0 is always an outlier. In fact, for a small number of traits under some conditions in this study (Supplemental table 3), and for root traits in other studies, it has been described to be rather average (e.g. Rosas et al. 2013).

### Genome-wide Association (GWA) Mapping and candidate genes

While phytohormone signaling pathways are fundamental mechanisms that are highly relevant for plant growth and development, only a few, small-scale studies have addressed them in the light of natural variation (Delker et al., 2010, Van Leeuwen et al., 2007). Consequently, these studies were not able to map any of the causal variants underlying the remarkable natural variation of responses to hormones. Using our data, we were able to search for common genetic variants that associate with altered hormone responses using GWAS. Using the large number of associations with ABA responses, we found that genes already implicated in the ABA pathway were not enriched, and possibly even underrepresented, in our set of GWAS candidate genes (Supplemental Figure 6). This supports the notion that genes that are targeted by natural variation are frequently not the genes that are found by traditional means (such as forward genetic mutant screens).

As more than 100 significant associations were detected, there were many good candidate genes and we needed to focus on the most interesting ones. We chose to study two cases with known candidate genes in regions associated with root traits upon hormone treatment. These turned out to be a interesting cases with regard to candidate gene approaches. The first case was an association for LRR that was detected under both control and CK treatment. We chose to investigate this region because a gene, *JAI3* that had been described to be involved JA signaling (Chini et al., 2007) was located within 20KB of the associated SNP (Figure 5B). As many hormonal signaling pathways are tightly intertwined (Vanstraelen & Benkova, 2012), it was tempting to assume *JAI3* was also involved in CK signaling. However, based on LD patterns, other genes were even better candidates, leading us to treat *JAI3* as one of 5 candidate genes in this region. Indeed, much like the LD pattern, gene expression analysis suggested that three other genes, and not *JAI3*, were the prime candidates in this region, based on the correlation of their expression pattern with the LRR trait (Figure 5C, Supplemental Figure 8B). The root traits observed in the available T-DNA insertion lines suggest that while *JAI3* has an effect on root length, this is not impacted by CK treatment and it is therefore not a likely candidate gene. In contrast, at least two other genes, At3g17890 encoding an unknown protein and At3g17910 encoding a protein similar to a cytochrome c oxidase assembly factor in human, are within the associated region and regulate root traits in response of CK (Figure 5D and 5E). These genes (and a third gene for which we could not obtain insertion lines) were the prime candidates based on LD and expression correlation with the trait and amongst each other (Supplemental Figure 8B). Consistently, both available lines showed an altered expression level of multiple hormone related genes. While cytokinin related genes were among those (most notably ARR12), in particular *PIN1* expression was altered strongly (Supplemental Figure 8D) in both lines and several other genes involved in the *SHY2* network were altered in the second mutant line (At3g17910). However, without further studies, it is difficult to clearly attribute the causality for these expression changes to a perturbation of hormone signaling or altered root morphology in these lines. Nevertheless, our data demonstrate that the yet uncharacterized genes At3g17890 and At3g17910 are involved in cytokinin dependent root growth regulation. This shows that an annotation based candidate gene approach can be misleading and that data-driven approaches (e.g. LD, expression) seem to be more efficient in identifying genes with a function in the GWAS trait of interest.

### *CRF3* is an integrator of multiple hormonal pathways

While further analysis of *JAI3* revealed that this is almost certainly not the causal gene, our second example showed that known genes should not be disfavored for consideration as causative candidates. In this second case, *CRF3*, a gene implicated in cytokinin and auxin signaling (Rashotte et al., 2006, Simaskova et al., 2015), was the sole candidate gene based on both LD and the absence of other genes in the region (Figure 6A and 6B). A published T-DNA insertion line (Simaskova et al., 2015) that led to loss of *CRF3* function displayed an ABA dependent alteration of root and lateral root traits (Figure 6C and 6D). At the molecular level, we observed a 3 to 4.5 fold increase in expression of *PIN1*, *PIN3*, and *PIN7* in whole roots of the *crf3* line, as well as altered expression of other genes involved in hormonal pathways (Supplemental Figure 8B). An opposite effect on *PIN1* and *PIN7* had been previously observed in root tips in this same line (Simaskova et al., 2015), suggesting that the effect of *CRF3* strongly depends on the tissue type (we used whole roots for our transcript measurements). Overall, our results suggest the possible involvement of *CRF3* as an integral part of 3 major hormone pathways, auxin and cytokinin, which were already known (Simaskova et al., 2015), and abscisic acid, as revealed by our study. Moreover, *CRF3* could be an integrator of hormonal and environmental stimuli, as it was recently shown that *CRF2* and *CRF3* regulate lateral root development under cold stress (Jeon et al., 2016).

### Local Adaptation and Hormone Signaling

Root traits are highly relevant for plant survival and productivity (Den Herder et al., 2010, Lynch, 1995). Hormonal pathways are major factors in the regulation of root traits and root system architecture (RSA). Here we showed that perturbation of hormonal pathways can shift RSA through the phenotypic space but the extent of this regulation can strongly vary between natural accessions of *Arabidopsis thaliana* (Fig. 4). As these accessions had been collected from different parts of the world, ranging from northern Sweden to southern Spain (Supplemental figure 1), we tested whether the variation in the root growth responses to hormones is associated with the climate at the places from which the accessions were derived (Supplemental table 9). Consistent with an important role of hormonal pathways for local adaptation, ABA regulated lateral root density and soil moisture in the coldest quarter of the year were the most important overlapping traits regarding the overlap of SNPs. The data indicate that less lateral roots are produced in an ABA dependent manner in accessions that are derived from areas in which the soil contains high moisture in the coldest quarter of the year. Support for the relevance of ABA related genes for local adaptation comes from studies on signatures of selection of ABA related genes when comparing closely related tomato species that grow in environments with contrasting precipitation regimes (Fischer et al., 2011, Xia et al., 2010) Moreover, in a study in *Arabidopsis thaliana*, ABA-responsive cis-regulatory elements (ABRE) showed significantly higher frequencies of polymorphisms in genes that responded to drought in a genotype dependent fashion than in drought responsive genes that responded similarly in all genotypes, suggesting that natural genetic variation in ABA responsive transcriptional control contributes to genotype specific drought responses (Lasky et al., 2014).

Despite 75% of all traits in our data were measured upon hormonal treatments, the majority of significant overlaps at the level of SNPs that were supported by significant trait correlations did not associate with hormonal treatments but with RSA traits under non-treated conditions. What does this mean for the importance of hormonal pathways for local adaptation? It is difficult to come up with a simple answer for this, as, also under control conditions, hormonal signaling pathways are involved in shaping root traits. Certainly, in the case of the link between ABA and Bio35, the data support an outstanding role for ABA for adapting to different soil moisture regimes in the coldest quarter, as no control condition for the same root trait is significantly associated with Bio35. The same holds true for TRL upon auxin treatment and Bio24 (Radiation of wettest quarter [W/m^2^]), as well as for P2 upon cytokinin treatment and Bio02 (Mean diurnal temperature range (mean(period max-min)) [°C]). Overall, we can conclude that natural variation in root traits is significantly associated with parameters that are highly relevant for local adaption in *Arabidopsis thaliana* and that, in particular, ABA regulated lateral root traits seem to be relevant for adaptation to soil moisture. Finally, we note that both SNP overlaps and trait correlations clearly indicate that we are far from explaining a major proportion of adaptation to climate variables, and that we expect numerous other processes and pathways to contribute to local adaptation. We would hope that similar systematic studies in the future will lead to a more complete picture that can then be tested in the wild and which might yield highly valuable insights into which traits and which underlying pathways contribute to local adaptation.

## MATERIALS & METHODS

### Plant Material and Growth Conditions

192 *Arabidopsis thaliana* accessions were used form the Busch group collection at GMI, Vienna. Names and numbers of all accessions are listed in Supplemental Table 1. Seeds were surface sterilized in a desiccator using 100 mL of household bleach and 3.5 mL of 37% HCl for 1 h. The seeds were stratified in water at 4°C in the dark for 3 days and were sown on square petri dishes containing 57 mL of growth medium. The control growth medium (before transfer) consisted of one-fifth-strength MS medium with MES buffer (Ducheva Biochemie), 1% Sucrose, 0.8% agar (Ducheva Biochemie), and adjusted to pH=5.7. Plates were dried for 45 min in a sterile laminar flow, before closing the cover and storing them at room temperature for three days before plating the seeds. In each plate, four accessions were plated, two in each row, with five plant replicates per accession (Supplemental Figure 2). Plates were placed vertically, and seeds were germinated under long-day conditions (21°C, 16-h-light/8-h-dark cycle). Individual seven-day-old seedlings were carefully transferred to the same position onto plates containing control medium supplemented with ABA, abscisic acid (1μM), CK, kinetin (1μM) or IAA, indole-acetic acid (0.5 μM) as indicated per experiment (Supplemental Figure 2).

Before performing the experiment on all 192 accessions, we first tested the effect of different MS (Murashige and Skoog) concentrations, since full-strength MS medium was found to inhibit root growth (Dubrovsky et al., 2009) and was originally designed for tissue culture that includes very high concentrations of mineral nutrients (Dubrovsky & Forde, 2012). We indeed observed inhibition of root growth in the tested concentrations (Supplemental figure 9). Thus, we decided to use the lowest concentration tested. Next, we tested the effect of placement in the upper vs. lower row in the plate, because we intended to fit 4 accessions on one plate. Thus, it was necessary to test if there is plate-position effect leading to differences in root growth between the two rows. Our results show that there was no significant difference between the rows in the plate in the tested accessions/concentrations (Supplemental Figure 10).

### Phenotypic Analysis

After three days on treatment, plates were scanned with CCD flatbed scanners (EPSON Perfection V600 Photo, Seiko Epson CO., Nagano, Japan) and images used to quantify root parameters with FIJI (Schindelin et al., 2012) (Supplemental Figure 2). We quantified: primary root length on day 10 (P), outgrowth of P on the treatment (P2), branching zone or the length of P between first and last visible lateral root (R), average lateral root length (LRL), and visible lateral root number (LR.No). Length was measured in millimeters (mm). Other traits were calculated: lateral root density of P, LRD_P (LR.No/P), lateral root density of R, LRD_R (LR.No/R), total lateral root length, TLRL (LR.No*LRL), total root length, TRL (TLRL+P), and length ratio, LRR (TLRL/P). For detailed protocol see (Ristova & Busch, 2017)

### Genome Wide Association Studies

We conducted GWA mapping on the mean trait values (in case of our root traits) and for each of the 35 climate variables capturing biologically relevant climate aspects (Bioclim variables) (Hutchinson et al., 2009, Kriticos et al., 2012) using a mixed model algorithm (Kang et al., 2008) which has been shown to correct for population structure confounding (Seren et al., 2012) and SNP data from the RegMap panel (Horton et al., 2012). SNPs with minor allele counts greater or equal to 10 were taken into account. The significance of SNP associations was determined at 5% FDR threshold computed by the Benjamini-Hochberg-Yekutieli method to correct for multiple testing (Benjamini & Yekutieli, 2001)

### Analyzing the overlap of ABA *bona fide* genes and ABA GWAS candidates

To determine a list of ABA *bona fide* candidate genes, we used the GO SLIM annotation of the TAIR10 release (ftp://ftp.arabidopsis.org/Ontologies/GeneOntology/ATHGOGOSLIM.txt; creation date: 4/9/15). We then filtered for GO annotations that contained the key word “abscisic acid” and were based on either IDA (“inferred by direct assay”) or IMP (“inferred by mutant phenotype”). The resulting list (Supplemental table 8) was subsequently used as the ABA *bona fide* candidate gene list. We then used the hypergeometric distribution to compute the P-value for the observed overlap of the GWAS based ABA candidate gene list and the as ABA *bona fide* candidate gene list. This was done in R (V3.1.2) using the command “dhyper(1,361,33602- 361,276)” where 1 is the overlap of genes between the two lists, 361 is the number of genes in the genome that were found in the GWAS, 33602-361 is the number of genes in the genome not found in the GWAS, and 276 is the number of genes in the genome that were *bona fide* candidate genes for ABA signaling.

### Computing the overlap of signals between GWASs

To determine whether the top associations between hormone GWAS and GWAS for environmental variables was higher than expected by chance, we first obtained the 0.5% most significantly associated SNPs in each GWAS. To avoid biases caused by outlier accessions, only SNPs with a minor allele count equal to or greater than 6 were included in the list of SNPs from which the top SNPs were derived. For each possible combination of root trait and climate variable GWAS, we computed the overlap of these top SNPs. We then used the hypergeometric distribution to compute a P-value for the observed overlap. This was done in python using the scipy.stats.hypergeom function. The list of all computed P-values for all possible combinations was corrected for multiple testing using the p.adjust function in R applying the “fdr” correction (Benjamini & Hochberg, 1995).

### Computing trait correlations

For computing the trait correlation, we considered only significant combinations of root and climate traits that we had identified using the SNP overlap analysis. We took only these lists into account as we only had a clear hypothesis for them, and testing all possible combinations without a clear hypothesis would have most likely resulted in an overly conservative penalization of P-values by multiple testing corrections for the very large number of tests that would have needed to be conducted. To calculate a population corrected association of root trait values and climate variables, we used the lm function of R and fitted a linear model containing either the first two eigenvectors that were obtained by performing a PCA on the SNP matrix from the RegMap population (Horton et al., 2012) (reduced model) or a model to which we added the climate variable (full model). We then performed an ANOVA to test whether the additional variable (climate) accounted for a significant proportion of variance.

### Statistical Analysis of Root Traits

A biological replicate for phenotyping purposes is an independent seedling of the same genotype. For mutant trait quantification, each assay was repeated 3 times, with each repetition having at least 10 plants per line and per conditions (Figure 5D,E, and Figure 6C,D). For RT-qPCR experiments, a biological replicate consisted of bulked roots grown on the same plate and technical replicates were cDNA samples that were split-up before RT-qPCR and independently measured. RT-qPCR was done with at least 3-4 biological and 2 technical replicates.

Statistical analysis and figure generation was done using JMP12 (SAS) (individual license to DR). Hierarchical clustering was performed using the average linkage method, using standardized mean values of the 10 root traits across the accessions and hormone treatments. The smallest number of clusters for each condition was chosen based on the distance graph (JMP12). Principal component analysis was performed on mean values of the 10 root traits across all accessions and conditions.

### Gene Expression Analysis (RT-PCR)

Total RNA extraction was performed using a commercial RNA isolation kit (RNeasy Mini Kit Plus, QIAGEN) using whole roots of 10-day-old seedlings, bulking 7-10 roots from the same accession/condition into one biological replicate. A qRT-PCR reaction was prepared using 2x SensiMix^TM^ SYBR & Fluorescein Kit (PEQLAB LLC, Wilmington, DE, USA) and PCR was conducted with a Roche Lightcycler^®^ 96 (Roche) instrument. Relative quantifications were performed for all the genes and the β-tubulin gene (AT5G62690) was used as an internal reference. Gene specific primers used are listed in Supplemental Table 11. Previously described primer pairs were used for *ARR1*, and *ARR12* (Moubayidin et al., 2010), *PIN3* and *PIN7* (Dello Ioio et al., 2008), and *TAA1* (Cui et al., 2013).

### Accession Numbers

Accession numbers for all 192 accessions of *Arabidopsis thaliana* used in this study can be found in Supplemental Table 1. T-DNA mutant lines used in this study: SALK_045920 (AT3G17910), SALK_050958 (AT3G17890), and SAIL_240_H09 (AT5G53290).

## ACKNOWLEDGMENTS

We thank Roberto Solano and Andrea Chini (Centro Nacional de Biotecnología, Spain) for kindly providing *jai3* seeds, and Matt Horton (University of Zurich) for providing the PCA Eigenvectors of the RegMap panel and for critical comments on the manuscript. We thank Samantha Krasnodebski for her help with media preparation, Christian Göschl for assistance in computing and programing, and Matt Watson for manuscript editing. Funding of this work was supported by the Austrian Academy of Science through the Gregor Mendel Institute (to D.R. and W.B.) and the European Union (FP7 COFUND PLANT FELLOWS Grant to D.R.).

### Author Contribution

DR, Conception and design, Contributed unpublished essential data or reagents, Acquisition of data, Analysis and interpretation of data, Drafting or revising the article; KM, Acquisition of data; WB, Conception and design, Contributed unpublished essential data or reagents, Analysis and interpretation of data, Drafting or revising the article.

## REFERENCES

Ackerly DD, Dudley SA, Sultan SE, et al., 2000. The evolution of plant ecophysiological traits: recent advances and future directions new research addresses natural selection, genetic constraints, and the adaptive evolution of plant ecophysiological traits. In: Bioscience, Pp.979–995., ed.

Alonso-Blanco C, Koornneef M, 2000. Naturally occurring variation in Arabidopsis: an underexploited resource for plant genetics. Trends Plant Sci 5, 22–9.

Atwell S, Huang YS, Vilhjalmsson BJ, et al., 2010. Genome-wide association study of 107 phenotypes in Arabidopsis thaliana inbred lines. Nature 465, 627–31.

Benjamini YA, Hochberg Y, 1995. Controlling the false discovery rate: a practical and powerful approach to multiple testing. Journal of the royal statistical society Series B (Methodological), pp.289–300.

Benjamini YA, Yekutieli D, 2001. The control of the false discovery rate in multiple testing under dependency. In: Annals of Statistics P-, ed.

Brachi B, Faure N, Horton M, et al., 2010. Linkage and association mapping of Arabidopsis thaliana flowering time in nature. PLoS Genet 6, e1000940.

Chini A, Fonseca S, Fernandez G, et al., 2007. The JAZ family of repressors is the missing link in jasmonate signalling. Nature 448, 666–71.

Cui D, Zhao J, Jing Y, et al., 2013. The arabidopsis IDD14, IDD15, and IDD16 cooperatively regulate lateral organ morphogenesis and gravitropism by promoting auxin biosynthesis and transport. PLoS Genet 9, e1003759.

De Smet I, Signora L, Beeckman T, Inze D, Foyer CH, Zhang H, 2003. An abscisic acid-sensitive checkpoint in lateral root development of Arabidopsis. Plant J 33, 543–55.

Delker C, Poschl Y, Raschke A, et al., 2010. Natural variation of transcriptional auxin response networks in Arabidopsis thaliana. Plant Cell 22, 2184–200.

Dello Ioio R, Nakamura K, Moubayidin L, et al., 2008. A Genetic Framework for the Control of Cell Division and Differentiation in the Root Meristem. Science 322, 1380–4.

Den Herder G, Van Isterdael G, Beeckman T, De Smet I, 2010. The roots of a new green revolution. Trends in Plant Science 15, 600–7.

Ding ZJ, Yan JY, Li CX, Li GX, Wu YR, Zheng SJ, 2015. Transcription factor WRKY46 modulates the development of Arabidopsis lateral roots in osmotic/salt stress conditions via regulation of ABA signaling and auxin homeostasis. Plant J 84, 56–69.

Du H, Wu N, Fu J, et al., 2012. A GH3 family member, OsGH3-2, modulates auxin and abscisic acid levels and differentially affects drought and cold tolerance in rice. Journal of Experimental Botany 63, 6467–80.

Dubrovsky JG, Forde BG, 2012. Quantitative analysis of lateral root development: pitfalls and how to avoid them. Plant Cell 24, 4–14.

Dubrovsky JG, Soukup A, Napsucialy-Mendivil S, Jeknic Z, Ivanchenko MG, 2009. The lateral root initiation index: an integrative measure of primordium formation. Ann Bot 103, 807–17.

Evans ML, Ishikawa H, Estelle MA, 1994. Responses of Arabidopsis Roots to Auxin Studied with High Temporal Resolution - Comparison of Wild-Type and Auxin-Response Mutants. Planta 194, 215–22.

Fischer I, Camus-Kulandaivelu L, Allal F, Stephan W, 2011. Adaptation to drought in two wild tomato species: the evolution of the Asr gene family. New Phytol 190, 1032–44.

Fournier-Level A, Korte A, Cooper MD, Nordborg M, Schmitt J, Wilczek AM, 2011. A map of local adaptation in Arabidopsis thaliana. Science 334, 86–9.

Gifford ML, Banta JA, Katari MS, et al., 2013. Plasticity Regulators Modulate Specific Root Traits in Discrete Nitrogen Environments. Plos Genetics 9.

Hong JH, Seah SW, Xu J, 2013. The root of ABA action in environmental stress response. Plant Cell Reports 32, 971–83.

Horton MW, Hancock AM, Huang YS, et al., 2012. Genome-wide patterns of genetic variation in worldwide Arabidopsis thaliana accessions from the RegMap panel. Nat Genet 44, 212–6.

Hu YF, Zhou G, Na XF, et al., 2013. Cadmium interferes with maintenance of auxin homeostasis in Arabidopsis seedlings. J Plant Physiol 170, 965–75.

Hutchinson M, T. Xu, D. Houlder, H. Nix A, Mcmahon J, 2009. ANUCLIM 6.0 User’s Guide Canberra: Australian National University, Fenner School of Environment and Society.

Kang HM, Zaitlen NA, Wade CM, et al., 2008. Efficient control of population structure in model organism association mapping. Genetics 178, 1709–23.

Kang JY, Choi HI, Im MY, Kim SY, 2002. Arabidopsis basic leucine zipper proteins that mediate stress-responsive abscisic acid signaling. Plant Cell 14, 343–57.

Koornneef M, Alonso-Blanco C, Vreugdenhil D, 2004. Naturally occurring genetic variation in Arabidopsis thaliana. Annu Rev Plant Biol 55, 141–72.

Kriticos DJ, Webber BL, Leriche A, et al., 2012. CliMond: global high-resolution historical and future scenario climate surfaces for bioclimatic modelling. Methods in Ecology and Evolution, 3(1), pp. 53–64.

Krouk G, Lacombe B, Bielach A, et al., 2010. Nitrate-Regulated Auxin Transport by NRT1.1 Defines a Mechanism for Nutrient Sensing in Plants. Developmental Cell 18, 927–37.

Kumar MN, Verslues PE, 2015. Stress physiology functions of the Arabidopsis histidine kinase cytokinin receptors. Physiol Plant 154, 369–80.

Laplaze L, Benkova E, Casimiro I, et al., 2007. Cytokinins act directly on lateral root founder cells to inhibit root initiation. Plant Cell 19, 3889–900.

Lasky JR, Des Marais DL, Lowry DB, et al., 2014. Natural variation in abiotic stress responsive gene expression and local adaptation to climate in Arabidopsis thaliana. Mol Biol Evol 31, 2283–96.

Leung J, Merlot S, Giraudat J, 1997. The Arabidopsis ABSCISIC ACID-INSENSITIVE2 (ABI2) and ABI1 genes encode homologous protein phosphatases 2C involved in abscisic acid signal transduction. Plant Cell 9, 759–71.

Li X, Mo X, Shou H, Wu P, 2006. Cytokinin-mediated cell cycling arrest of pericycle founder cells in lateral root initiation of Arabidopsis. Plant Cell Physiol 47, 1112–23.

Liu W, Li RJ, Han TT, Cai W, Fu ZW, Lu YT, 2015. Salt Stress Reduces Root Meristem Size by Nitric Oxide-Mediated Modulation of Auxin Accumulation and Signaling in Arabidopsis. Plant Physiology 168, 343–U607.

Lopez-Bucio J, Hernandez-Abreu E, Sanchez-Calderon L, et al., 2005. An auxin transport independent pathway is involved in phosphate stress-induced root architectural alterations in arabidopsis. Identification of BIG as a mediator of auxin in pericycle cell activation. Plant Physiology 137, 681–91.

Lynch J, 1995. Root Architecture and Plant Productivity. Plant Physiology 109, 7–13.

Malamy JE, 2005. Intrinsic and environmental response pathways that regulate root system architecture. Plant Cell and Environment 28, 67–77.

Moubayidin L, Perilli S, Dello Ioio R, Di Mambro R, Costantino P, Sabatini S, 2010. The rate of cell differentiation controls the Arabidopsis root meristem growth phase. Curr Biol 20, 1138–43.

Mouchel CF, Briggs GC, Hardtke CS, 2004. Natural genetic variation in Arabidopsis identifies BREVIS RADIX, a novel regulator of cell proliferation and elongation in the root. Genes Dev 18, 700–14.

Mouchel CF, Osmont KS, Hardtke CS, 2006. BRX mediates feedback between brassinosteroid levels and auxin signalling in root growth. Nature 443, 458–61.

Nacry P, Canivenc G, Muller B, et al., 2005. A role for auxin redistribution in the responses of the root system architecture to phosphate starvation in Arabidopsis. Plant Physiol 138, 2061–74.

Pacheco-Villalobos D, Sankar M, Ljung K, Hardtke CS, 2013. Disturbed local auxin homeostasis enhances cellular anisotropy and reveals alternative wiring of auxin-ethylene crosstalk in Brachypodium distachyon seminal roots. PLoS Genet 9, e1003564.

Perez-Torres CA, Lopez-Bucio J, Cruz-Ramirez A, et al., 2008. Phosphate availability alters lateral root development in Arabidopsis by modulating auxin sensitivity via a mechanism involving the TIR1 auxin receptor. Plant Cell 20, 3258–72.

Rashotte AM, Mason MG, Hutchison CE, Ferreira FJ, Schaller GE, Kieber JJ, 2006. A subset of Arabidopsis AP2 transcription factors mediates cytokinin responses in concert with a two-component pathway. Proc Natl Acad Sci U S A 103, 11081–5.

Ristova D, Busch W, 2017. Genome-Wide Association Mapping of Root Traits in the Context of Plant Hormone Research. Methods Mol Biol 1497, 47–55.

Ristova D, Rosas U, Krouk G, Ruffel S, Birnbaum KD, Coruzzi GM, 2013. RootScape: a landmark-based system for rapid screening of root architecture in Arabidopsis. Plant Physiol 161, 1086–96.

Rosas U, Cibrian-Jaramillo A, Ristova D, et al., 2013. Integration of responses within and across Arabidopsis natural accessions uncovers loci controlling root systems architecture. Proc Natl Acad Sci U S A 110, 15133–8.

Ruffel S, Krouk G, Ristova D, Shasha D, Birnbaum KD, Coruzzi GM, 2011. Nitrogen economics of root foraging: Transitive closure of the nitrate-cytokinin relay and distinct systemic signaling for N supply vs. demand. Proceedings of the National Academy of Sciences of the United States of America 108, 18524–9.

Salome PA, To JPC, Kieber JJ, Mcclung CR, 2006. Arabidopsis response regulators ARR3 and ARR4 play cytokinin-independent roles in the control of circadian period. Plant Cell 18, 55–69.

Satbhai SB, Ristova D, Busch W, 2015. Underground tuning: quantitative regulation of root growth. Journal of Experimental Botany 66, 1099–112.

Schindelin J, Arganda-Carreras I, Frise E, et al., 2012. Fiji: an open-source platform for biological-image analysis. Nat Methods 9, 676–82.

Seren U, Vilhjalmsson BJ, Horton MW, et al., 2012. GWAPP: a web application for genome-wide association mapping in Arabidopsis. Plant Cell 24, 4793–805.

Simaskova M, O’brien JA, Khan M, et al., 2015. Cytokinin response factors regulate PIN-FORMED auxin transporters. Nat Commun 6, 8717.

Singh AP, Fridman Y, Friedlander-Shani L, Tarkowska D, Strnad M, Savaldi-Goldstein S, 2014. Activity of the Brassinosteroid Transcription Factors BRASSINAZOLE RESISTANT1 and BRASSINOSTEROID INSENSITIVE1-ETHYL METHANESULFONATESUPPRESSOR1/BRASSINAZOLE RESISTANT2 Blocks Developmental Reprogramming in Response to Low Phosphate Availability. Plant Physiology 166, 678–88.

Stepanova AN, Robertson-Hoyt J, Yun J, et al., 2008. TAA1-mediated auxin biosynthesis is essential for hormone crosstalk and plant development. Cell 133, 177–91.

Sun P, Tian QY, Chen J, Zhang WH, 2010. Aluminium-induced inhibition of root elongation in Arabidopsis is mediated by ethylene and auxin. Journal of Experimental Botany 61, 347–56.

Van Leeuwen H, Kliebenstein DJ, West MA, et al., 2007. Natural variation among Arabidopsis thaliana accessions for transcriptome response to exogenous salicylic acid. In: The Plant Cell, Pp.2099–2110., ed.

Vanstraelen M, Benkova E, 2012. Hormonal interactions in the regulation of plant development. Annu Rev Cell Dev Biol 28, 463–87.

Vicente-Agullo F, Rigas S, Desbrosses G, Dolan L, Hatzopoulos P, Grabov A, 2004. Potassium carrier TRH1 is required for auxin transport in Arabidopsis roots. Plant Journal 40, 523–35.

Vidal EA, Araus V, Lu C, et al., 2010. Nitrate-responsive miR393/AFB3 regulatory module controls root system architecture in Arabidopsis thaliana. Proc Natl Acad Sci U S A 107, 4477–82.

Werner T, Motyka V, Strnad M, Schmulling T, 2001. Regulation of plant growth by cytokinin. Proc Natl Acad Sci U S A 98,10487–92.

Xia H, Camus-Kulandaivelu L, Stephan W, Tellier A, Zhang Z, 2010. Nucleotide diversity patterns of local adaptation at drought-related candidate genes in wild tomatoes. Mol Ecol 19, 4144–54.

Yamada H, Suzuki T, Terada K, et al., 2001. The Arabidopsis AHK4 histidine kinase is a cytokinin-binding receptor that transduces cytokinin signals across the membrane. Plant and Cell Physiology 42, 1017–23.

Yu J, Pressoir G, Briggs WH, et al., 2006. A unified mixed-model method for association mapping that accounts for multiple levels of relatedness. Nat Genet 38, 203–8.

Yuan HM, Xu HH, Liu WC, Lu YT, 2013. Copper regulates primary root elongation through PIN1-mediated auxin redistribution. Plant Cell Physiol 54, 766–78.

Zhao YK, Wang T, Zhang WS, Li X, 2011. SOS3 mediates lateral root development under low salt stress through regulation of auxin redistribution and maxima in Arabidopsis. New Phytologist 189, 1122–34.

